# Investigating the Central Dogma of Molecular Biology in the Context of Nuclear Architecture and Cell Cycle

**DOI:** 10.1101/810499

**Authors:** Shivnarayan Dhuppar, Aprotim Mazumder

## Abstract

Nuclear architecture is the organization of the genome within a cell nucleus with respect to different nuclear landmarks such as nuclear lamina, matrix or nucleoli. Lately it has emerged as a major regulator of gene expression in mammalian cells. The studies connecting nuclear architecture with gene expression are largely population-averaged and do not report on the heterogeneity in genome organization or in gene expression within a population. In this report we present a method for combining 3D DNA Fluorescence *in situ* Hybridization (FISH) with single molecule RNA FISH (smFISH) and immunofluorescence to study nuclear architecture-dependent gene regulation on a cell-by-cell basis. We further combine it with an imaging-based cell cycle staging to correlate nuclear architecture with gene expression across the cell cycle. We present this in the context of *Cyclin A2 (CCNA2)* gene for its known cell cycle-dependent expression. We show that, across the cell cycle, the expression of a *CCNA2* gene copy is stochastic and depends neither on its sub-nuclear position—which usually lies close to nuclear lamina—nor on the expression from the other copies.

## Introduction

Life at the cellular level is the result of the coordinated efforts of molecular machinery inside a cell brought about by timely expression of genes at mRNA and protein levels. A cell achieves this by regulating the genes at various stages of their expression—from the transcription of the genes to the translation of the respective mRNAs and eventual degradation of the gene products [1–3]. Many of these effectors of gene regulation can be linked to two important factors: nuclear architecture and the cell cycle [4, 5]. Cell cycle is the complete gamut of processes with which a cell, starting as a single cell, divides into two daughter cells. This implies that almost all cellular events or attributes have to be coordinated with the cell cycle to ensure proper cell division in the end—even the nuclear architecture [6– 8]. Nuclear architecture is the way genome is organized inside a cell nucleus with respect to nuclear compartments such as the lamina or heterochromatin. It can play a key role in determining the gene expression pattern in a cell and hence the cellular identity as proposed by the topological model of gene regulation (TMGR) [9]. The TMGR hypothesizes two ideas: 1) spatial positioning of a gene with respect to chromosomal locations such as centromeres can affect the transcription of that gene and 2) differentiating cells develop a pattern of gene positions with respect to nuclear compartments such as nuclear lamina or heterochromatin or interchromatin compartments which defines the patterns of gene expression and hence the eventual identities of those cells [9]. There have been a number of studies supporting the TMGR but most of them lacked either the single-cell resolution or the ability to interrogate gene position and expression at the same time in the same cells [10–13]. Single cell level studies on nuclear architecture-dependent gene expression have been elusive as they require the combination of immunofluorescence-based detection of proteins with fluorescent *in situ* hybridization (FISH)-based detection of gene positions and mRNAs while also preserving the 3D nuclear architecture of the cell. The existing methods for RNA FISH and 3D DNA FISH are incompatible because the steps involved in either may affect the detection of the other adversely.

Here we present a simple and reliable way to combine DNA FISH with single molecule RNA FISH (smFISH) and immunofluorescence while also preserving the 3D nuclear architecture of a cell. We further link it with the microscopy-based cell cycle staging developed earlier [14] to study cell cycle-dependent changes in nuclear architecture and gene expression on a cell-by-cell basis. In particular, we investigate how the positioning of the *Cyclin A2* gene with respect to the nuclear lamina correlates with the already-known cell cycle-dependent expression of the gene.

The ease of the technique presented here combined with the depth of the analyses makes it a useful tool in addressing various questions pertaining to cell cycle, nuclear architecture and gene expression at a single-cell resolution.

## Results

### Combined DNA FISH, smFISH and immunofluorescence in 3-dimensionally intact nuclei

Genome organization has previously been shown to be involved in the regulation of gene expression. Most of these studies either rely on bulk biochemical assays for measuring the expression [15] or involve FISH-based labeling and imaging of the mRNA followed by the same for the gene of interest—this relies heavily on the ability to mark and identify the same cells between the two steps [11, 16, 17]. Recent years have seen the development of Single molecule RNA FISH (smFISH) methods that yield absolute transcript counts on a cell by cell basis unlike standard RNA FISH where only relative intensities are measured [18, 19]. While there have been previous studies describing combined DNA and RNA FISH [11, 16, 17] fewer attempts have been made to combine 3D DNA FISH with smFISH. The difficulty in combining DNA FISH with RNA FISH or immunofluorescence stems from the fact that DNA FISH involves harsh treatments such as acid-or formamide-based DNA denaturation, which can adversely interfere with the subsequent detection of proteins or RNAs inside a cell [10, 20]. Besides, the ethanol dehydration involved in such procedures destroys the 3D architecture of the cells—which are more than 50% water—by flattening them out. The organelle most affected by such dehydrations would be the cell nucleus whose water content can be as high as 85% [21]. This makes it difficult to combine the three assays while also keeping the distortions to nuclear architecture to minimum. Besides, the existing protocols for 3D DNA FISH involve steps such as nitrogen freeze-thaw cycles which add another layer of difficulty to the problem [22].

We started out by first standardizing a protocol for 3D DNA FISH devoid of conventional difficulties. We realized that the major factor, and perhaps the only one, affecting the 3D nuclear architecture of a cell in a conventional 2D DNA FISH protocol is ethanol dehydration. We discovered that the loss in the efficiency of probe hybridization due to omission of ethanol dehydration can be compensated by having longer hybridization times (> 40 hours). The resultant protocol, in fact, is largely similar to a conventional 3D DNA FISH protocol if nitrogen freeze-thaw cycles are removed.

Once the protocol for 3D DNA FISH was standardized, the next step was to determine the order in which the three techniques can be performed for a least interference among them. We observed that the order which gives the best result is the following: immunofluorescence followed by refixation with 4% paraformaldehyde followed by DNA FISH and then smFISH for RNA. The order is especially important with regard to smFISH where 20-nucleotide long oligomers are used as the hybridization probes. If not performed in the correct order, the high formamide concentration in DNA FISH hybridization buffers can destroy all smFISH RNA signals.

Once standardized we used the above technique to study how gene position inside the nucleus can be correlated with the gene expression across the cell cycle in HeLa cells. We selected the *Cyclin A2 (CCNA2)* gene for our study due to its known cell cycle-dependent gene expression. We found that most of the cells had three copies of *CCNA2* gene, which was consistent with the fact that HeLa cells are hypertriploid (Figure 1A). We also observed that mRNA and protein expression for *CCNA2* gene were strongly correlated (Figure 1A) and that the activity of two adjacent alleles, as inferred from the colocalization of the smFISH and DNA FISH signals, could be very different—at least in 2D-projected images of nuclei (Figure 1B).

**Figure 1:**
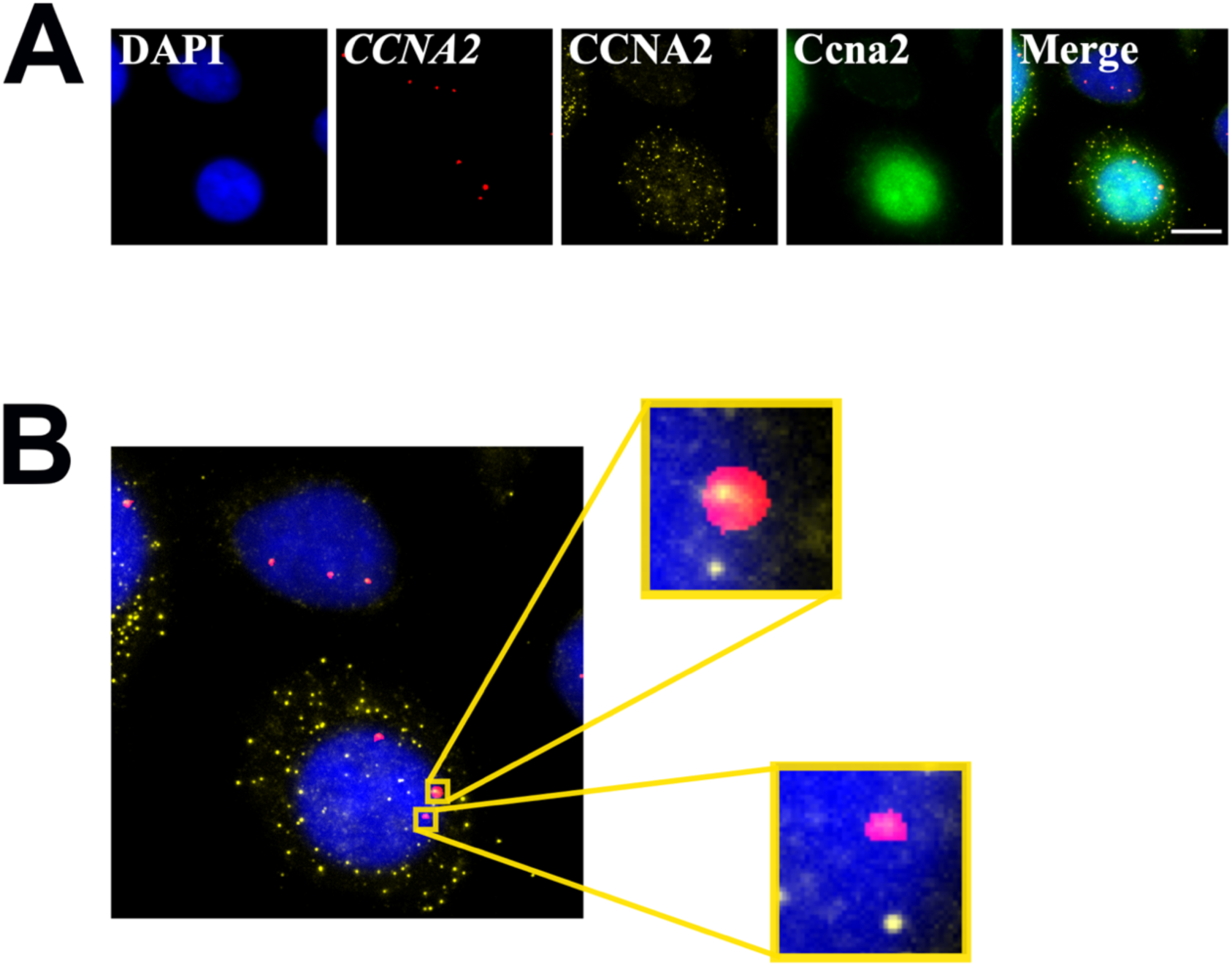
Combined DNA FISH, smFISH and immunofluorescence. (A) Combined DNA FISH, smFISH and immunofluorescence for *CCNA2* gene. The protein and mRNA expressions are correlated - *CCNA2* mRNA and protein are both low in the cell above, while they are high in the cell below. For a detailed quantification over many cells across cell cycle phases please see Figure 3E. (B) Two adjacent copies of *CCNA2* gene with different activity status. Nascent transcripts are seen at the gene locus above, but not below.

To convert the above observations to quantifiable, comparable metrics we performed a quantitative 3D image analysis as discussed next.

### 3D nuclear segmentation and cell cycle staging

The cells were first analyzed for their DNA content using a previously-developed module for imaging-based cell cycle staging [14]. Briefly the workflow ran thus: the average-projected images from the DAPI channel were passed to the module quantifying the DNA contents of individual nuclei in a field. Once DNA content was quantified the 3D stack for each nucleus in a field was sent one-by-one to another module which eliminated out-of-focus planes from the 3D stacks and segmented out the nucleus in 3D based on texture analysis (Figure 2A). All 3D analyses were done in a cell-by-cell manner to reduce the size of the images to what a desktop computer can easily process. DNA FISH spots were also segmented in 3D using a module again from our previous report [14]. Figure 1B shows an example of one such 3D segmented nucleus and the segmented *CCNA2* alleles within (see also Supplementary Video V1). Also the nuclear volumes obtained from the above analysis compared well with what has been reported previously for HeLa cells (Figure 1C) [23].

**Figure 2:**
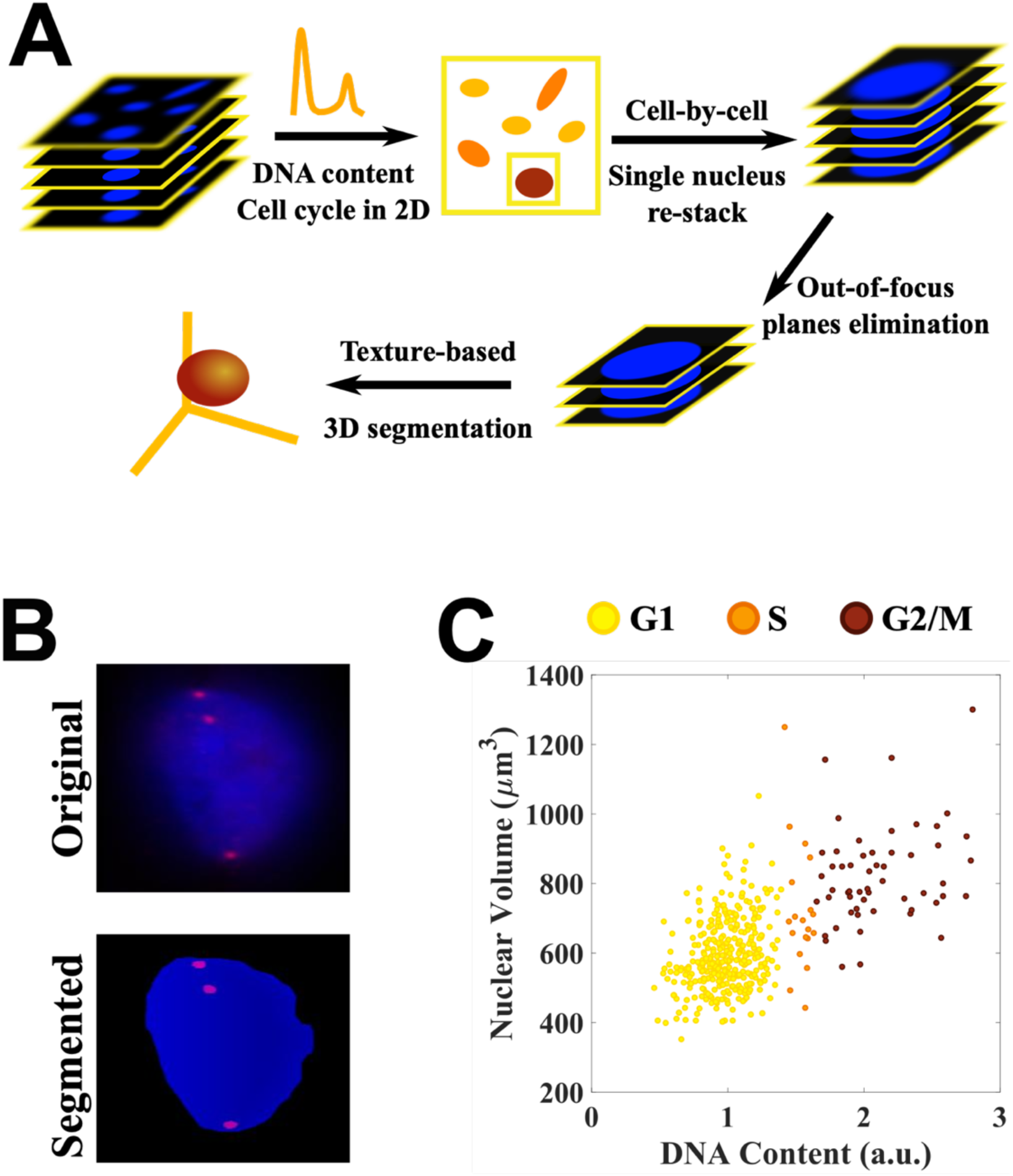
3D nuclear segmentation and cell cycle staging. (A) The workflow for 3D nuclear segmentation. Cells were first analyzed for their DNA content in 2D. Thereafter 3D stack for each nucleus was sent one-by-one to an image processing module which removed out-of-focus light and segmented out the individual nuclei in 3D using texture-based analysis. (B) A sample segmentation of a nucleus (blue) and the *CCNA2* gene copies within (red). (C) The volume measurements for HeLa cell nuclei from the above analysis match with the ones reported. See Supplementary Video V1 for visualization of the 3D segmentation shown in (B).

The 3D segmented nuclei and alleles thus obtained were then used to calculate distances of the centroid of an allele from the nuclear lamina and nuclear centroid to define a Closeness metric as discussed below.

### Central dogma in the context of nuclear architecture and cell cycle

The topological model of gene regulation suggests that upon differentiation cells match the expression patterns of their differentiated states by defining a unique reproducible pattern of gene positions with respect to different nuclear compartments such as the nuclear lamina, heterochromatin or interchromatin compartments [9]. Yet another theory suggests that genes can make long extrusions out of their chromosome territories for their transcription [10, 24]. This looping out of genes for the expression could be to avoid nuclear compartments such as nuclear lamina or heterochromatin which are generally associated with gene silencing [25– 27]. To determine which of the two hypotheses—the static TMGR model or the dynamic extrusion-based model—is true in the case of *CCNA2* gene, we calculated the distances of the centroids of the three copies from the nuclear lamina and nuclear centroid and then correlated them with the expression of *CCNA2* gene.

We observed that although the expression of *CCNA2* gene in terms of mRNA numbers and protein levels peaked in the G2/M phases of the cell cycle (Figures 3A and 3B), the means of the absolute distances of the centroids of the gene copies from nuclear lamina or the centroid did not change substantially across the cell cycle (Figures 3C and 3D). We further defined a Closeness metric for the gene copies such that its value always lies between 0 and 1—unity if the copy is at the lamina and zero if it is at the center. We numbered the gene copies such that copy 1 always corresponded to the highest Closeness followed by copy 2 and then copy 3. We observed that the mean value of Closeness in a cell did not correlate with the expression of *CCNA2* gene in terms of mRNA number or protein level (Figure 3E). In fact, the mean Closeness for the *CCNA2* gene did not vary much across the cell cycle even though the expression was highly cell cycle-dependent (Figure 3E).

**Figure 3:**
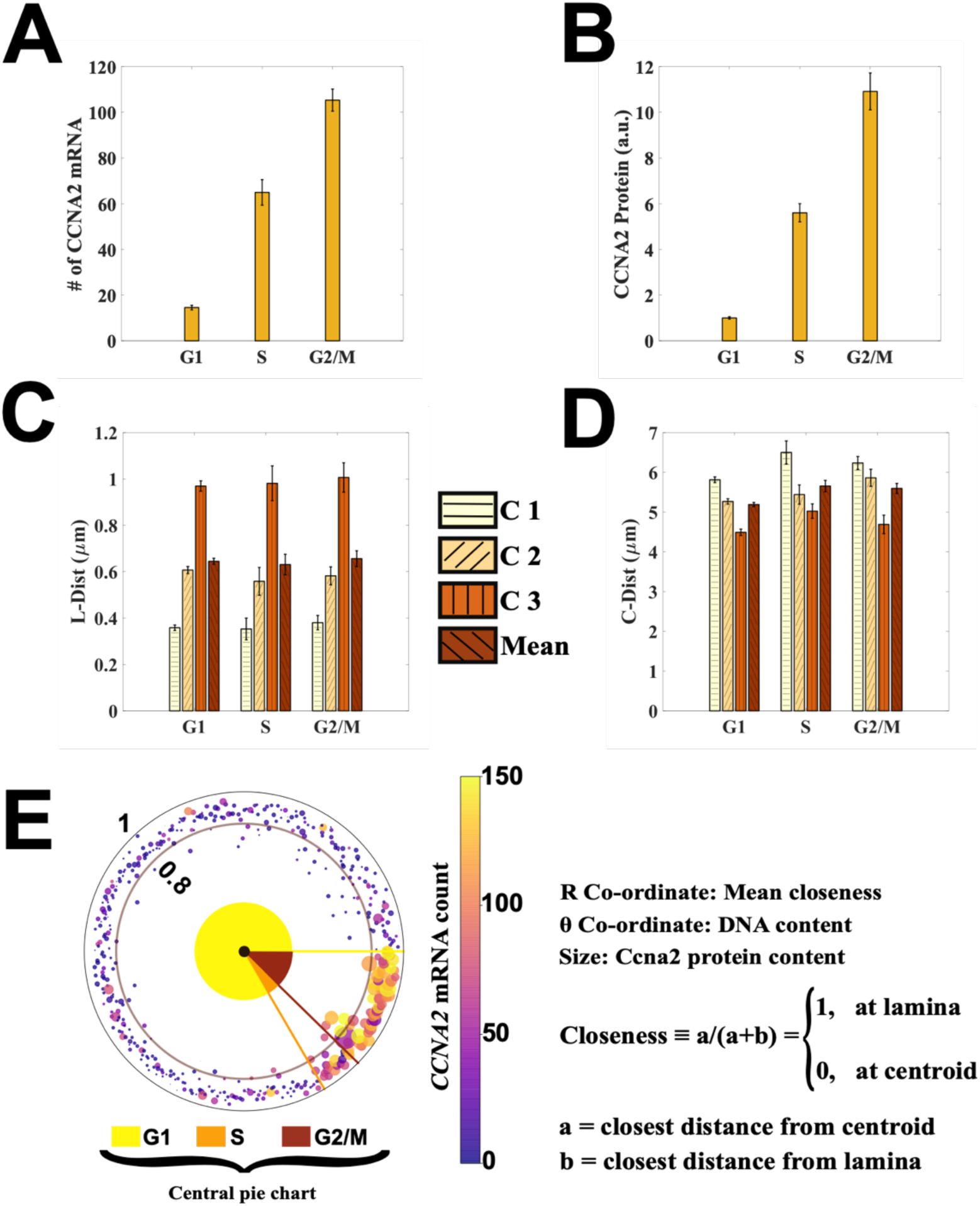
Central dogma in the context of nuclear architecture and cell cycle. (A) Mean of *CCNA2* mRNA number peaks in the G2/M phase of the cell cycle. (B) Mean of *CCNA2* protein levels also peaks in the G2/M phase of the cell cycle. (C) Means of the distances of the three copies from the nuclear lamina in the three cell cycle phases. Copies are numbered according to their relative distances from the lamina (L-Dist) with C1 (Copy 1) closest to the lamina followed by C2 and then C3. (D) Means of distances of the three copies from the centroid of the nucleus (C-dist). The means of the absolute values of both the distances do not change much across the cell cycle. (E) A polar plot representing how *CCNA2* expression is correlated with the average closeness of the three copies in a cell. Each spot in the graph is an individual cell, with spot color representing corresponding *CCNA2* mRNA count and spot size Ccna2 protein content. While mRNA and protein expressions for the *CCNA2* gene are well correlated across the cell cycle (both are high in S, G2 cells), there is no obvious dependence of *CCNA2* expression on the average closeness of its copies from the lamina. Pie chart in the center represents the cell cycle distribution of the population.

The above observations strongly support TMGR mode of gene regulation in the case of *CCNA2* gene in the differentiated cancer cell line. But the analysis is incomplete without the information on the activity of the gene at a single-allele level. The next result discusses the position-dependent activity of *CCNA2* gene at a single-allele level.

### Position-independent expression of *CCNA2* gene copies across the cell cycle

We determined the activity of a gene copy by finding the overlap between smFISH and DNA FISH signals at a position in the nucleus in 3D. A cuboid just enveloping the gene spot was constructed which then was searched for the mRNA signal. If the cuboid contained at least an mRNA spot then that gene copy was deemed as actively expressing.

We observed that there was no obvious correlation between the position of a *CCNA2* allele and its expression across the cell cycle. As shown in Figure 4, the expression of a copy did not depend on its position relative to the lamina or the cell cycle stage: it could express in any cell cycle stage irrespective of its subnuclear position—though the overall mRNA levels are cell cycle dependent. We also investigated the number of *CCNA2* copies actively expressing in a cell and the corresponding total mRNA count. We found that the cells with higher mRNA counts, on the average, had more gene copies with active transcription (Supplementary Figure S1). But interestingly, we also found cells with very small mRNA counts too can have an active expression from more than a single copy and vice versa. This illustrates stochastic nature of gene activation via transcriptional bursts (Supplementary Figure S1) [28].

**Figure 4:**
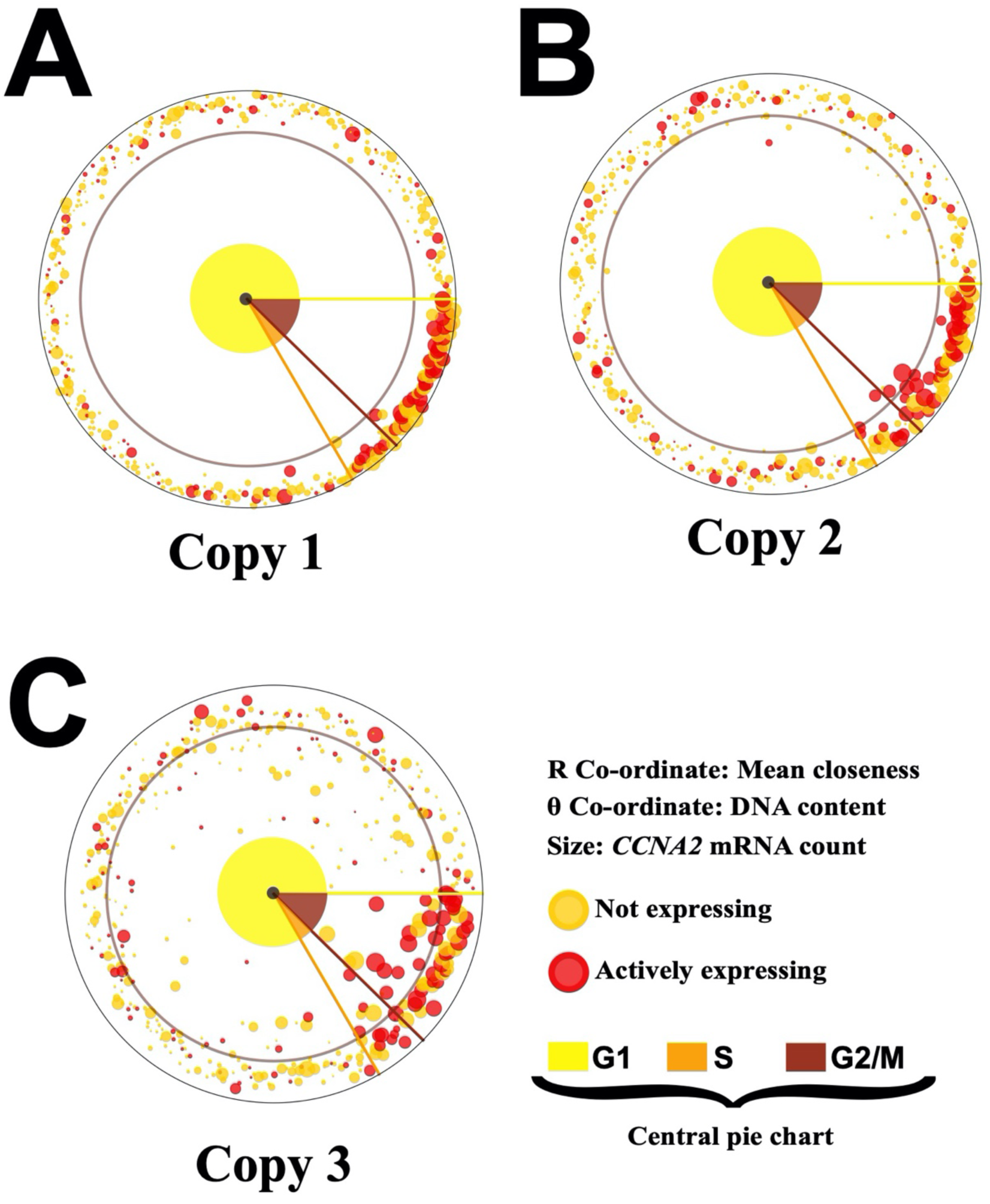
Position-independent transcription of the *CCNA2* gene copies across the cell cycle. (A, B and C) Polar plots representing the position-dependent activity of the three copies individually in a cell across the cell cycle. Copies are numbered such that Copy 1 is closest to the lamina followed by Copy 2 and then Copy 3. There is no correlation between the activity of a copy and its relative distance from the lamina, though a copy is more likely to express in the S or G2/M phases of the cell cycle. See also Supplementary Figure S1 for the cell cycle distribution of the number of *CCNA2* gene copies actively expressing in a cell.

We also observed that two of the three alleles always stayed close to the nuclear lamina (Figure 4)—this is striking given that individual bona fide lamina-associated domains (LADs) can get randomized between two cell cycles [29] even if the overall nuclear organization of chromosome territories can remain the same [30].

The above observations support the TMGR mode of action for *CCNA2* gene regulation where the average position of the gene relative to nuclear lamina varies little within the cell cycle even though its expression—both at mRNA and protein levels—is highly cell cycle-dependent.

### Discussion

Nuclear architecture derives from the non-random organization of the eukaryotic genome in terms of its arrangement of chromosome territories and genes with respect to one another and to other nuclear landmarks such as nuclear lamina or nucleoli [31]. It is involved in cell fate decisions during differentiation [10, 32, 33] and has been shown to be tissue-specific [34]. It is also implicated in the regulation of gene expression via association with different nuclear compartments such as nuclear periphery, heterochromatin, euchromatin or interchromatin compartments [35, 36]. Most of these studies are based on bulk biochemical assays which can efficiently capture population-level changes in organization or expression but miss out on the cell-to-cell variability within a population. In fact, according to recent reports there is a substantial level of heterogeneity in genome organization within a population and that at a time only a small fraction of cells within a population harbor the interactions captured at the population-level [8, 37]. This underscores the need for single cell studies to capture nuclear architecture-dependent regulation of DNA processes in their entirety.

The existing methods to study 3D nuclear architecture and the gene expression at a single-cell resolution are incompatible with one another. Here we report a simple and efficient way to combine DNA FISH with smFISH for RNA and immunofluorescence which also preserves the 3D nuclear architecture of a cell. We further combine this with an imaging-based cell cycle staging developed earlier [14] to study gene regulation in the context of nuclear architecture across the cell cycle for a cell cycle-regulated gene *Cyclin A2*.

We observe that the cell cycle dependence of *CCNA2* gene expression is hardly reflected in the average positioning of *CCNA2* gene relative to the nuclear periphery (Figure 3E). Interestingly in contrast to previous studies associating nuclear periphery to gene silencing [25–27], we observe that nuclear positioning of a *CCNA2* gene—which more often stays closer to the nuclear periphery—matters little for its activity (Figure 4). Also, the number of *CCNA2* copies actively expressing in a cell do not reflect on the overall mRNA count in that cell—there can be cells with very high mRNA count but fewer than two copies actively expressing and vice versa. This demonstrates the stochastic nature of transcriptional bursts and gene regulation via the modulation of burst frequency [38]. This might also suggest a possible decoupling between the activities of different alleles within a nucleus where each is agnostic of others’ transcriptional status unlike as reported recently for the bacterial system [39]. The above observations further emphasize the importance of studying the heterogeneity in genome organization within a population at a single cell resolution.

Taken together, the technique presented here along with the analysis modules makes it a powerful tool to study nuclear architecture or nuclear architecture-dependent gene regulation across the cell cycle at a single cell resolution.

## Materials and methods

### Cell culture

HeLa cells were grown on ethanol-sterilized, ploy-D-lysine-treated coverslips. DMEM (Gibco) supplemented with 10% FBS (Gibco) without any antibiotic was used for the cell culture. Cells were tested negative of any bacterial contamination including mycoplasma. They were passaged every ∼4 days and were let to grow for at least 24 hours before starting an experiment.

### Combined DNA FISH, smFISH and immunofluorescence

Cells were fixed with 4% paraformaldehyde (PFA) in nuclease-free (NF) PBS for 15 minutes at room temperature (RT) followed by permeabilization with 0.3% Triton-x 100 in NF PBS for 10 minutes at RT. The cells were then washed twice with PBS and were prepared for immunofluorescence. The order of assays was always thus: immunofluorescence followed by re-fixation for 15 minutes with 4% PFA in NF PBS at RT followed by DNA FISH followed by RNA FISH. The re-fixation post immunofluorescence is important to preserve the signals.

For immunofluorescence, cells were first blocked for non-specific binding using 1% NF BSA from Ambion for 30 minutes at RT. Then the cells were incubated with the primary antibodies in 1% NF BSA for 60 minutes at RT. After two washes with NF PBS the cells were incubated with the secondary antibodies in 1% NF BSA for 60 minutes at RT followed by two washes with NF PBS.

For DNA FISH, cells were first washed with 2X NF SSC. After which the DNA was denatured with 70% formamide in 2X NF SSC at 80 °C for 15 minutes. While DNA in the cells was denatured, DNA hybridization mixture was prepared: 9 µl hybridization buffer (from Empire Genomics) + 1 µl probes. The mixture was heated at 80°C for 3 minutes for denaturing the probes. A slide was wipe cleaned with RNase Zap and let to dry. After DNA denaturation, the DNA FISH hybridization mix was transferred on the slide and the coverslip was placed over it carefully. The cells were then incubated for hybridization for more than 40 hours at 37 °C in a dark heavily-humidified chamber. After hybridization the cells were washed once with 2X SSC at RT followed by two 10-miunte washes with 50% formamide in 2X NF SSC at 37 °C.

For smFISH the cells were first rinsed with 2X NF SSC followed by two quick washes with 10% formamide in 2X NF SSC at RT. smFISH hybridization mix was prepared thus: 9 µl Hybridization buffer (Stellaris) + 1 µl formamide + 0.5 µl RNA FISH probes. The slides were wiped clean with RNase Zap and were let to dry completely. The hybridization mix was then transferred onto the slide and the coverslip was carefully placed over it. The cells were then incubated overnight at 37 °C in a dark heavily-humidified chamber. The next day cells were washed once with 2X SSC at RT followed by two 5-minute washes with 10% formamide in 2X NF SSC at 37 °C.

Finally the cells were DNA stained with 1 µg/ml Hoechst 33342 dye for 10 minutes.

### Antibodies, FISH probes and chemicals used

Ccna2 protein was labelled with a rabbit antibody (1:500) from Abcam (ab181591). For detection goat anti-rabbit antibody (1:500) with Alexa Fluor 647 dye was used. Pre-labelled *CCNA2* DNA FISH probes were procured from Empire Genomics (CCNA2-20-Re). The fluorophore was spectrally similar to Alexa Fluor 594. It came with the hybridization buffer.

For RNA FISH the probes were designed using Stellaris Probe Designer by Biosearch technologies and were ordered from the same source. The probes were labeled with Quasar 570 fluorophores. The probes sequences are available in the Supplementary Table 1 and have been described before [14].

Ambion ultrapure 50g/l BSA (AM2616) was procured from Invitrogen.

All other reagents were Ambion RNase-free products from Invitrogen.

### Microscopy

Imaging was always performed with Vectashield as a mounting medium.

Images were acquired at 14-bit resolution using the fully-automated Olympus IX83 microscope on a Retiga 6000 camera (QImaging). All images were take using a 60X oil objective with 1.42 NA. 31 planes at 300 nm apart were taken for every field. Appropriate filter cubes from Chroma Technologies (49309 ET – Orange#2 FISH and 49310 ET – Red#2 FISH) and Olympus were used for preventing any bleed through across the spectrum of fluorophores.

### Image and data analysis

The 3D stacks were first average projected and analyzed for DNA content using an automated Matlab routine developed earlier [14]. Thereafter the 3D stack corresponding to each nucleus in the field was sent one-by-one to another module for texture-based removal of the out of focus light and finally 3D segmentation of the nucleus (Figure 2). After segmentation the boundary voxels and the centroid of each cell nucleus were used for finding relevant distances. A Closeness metric was defined for relative distances of the alleles from the lamina and the centroid thus: ratio of the distance of an allele from the centroid to the sum of the distances from the lamina and the centroid (Figure 3). Its value lies always between 0 and 1: unity if the allele is on the lamina and zero if it is at the centroid. The 3D segmentation was done on a cell-by-cell basis to reduce the size of the images and hence make the computation feasible with a desktop computer.

For position-dependent analysis of gene expression, only nuclei with exactly three clearly identified copies of *CCNA2* gene were used—they formed ∼73% of total population.

The physical size of a pixel on the camera was 4.54 µm corresponding to an area of 20.6 µm^2^. With a 60X, 1.42 NA objective without binning it translates to 5.73 × 10^−3^ µm^2^. With stack separation of 300 nm, the voxel volume comes to 1.72 × 10^−3^ µm^3^. Distances in 3D were found similarly.

All image analyses were performed on Matlab. For graphs Matlab and Python 3 were used. All codes and programs used in the study are available on Github.com/shuppar.

## Author contributions

AM, SD conceived and designed the study. SD performed the experiments and the analysis. AM, SD interpreted the data and wrote the manuscript. AM monitored the project and associated grants.

## Acknowledgements

SD thanks Dr Sitara Roy for many useful suggestions on data representation and visualization.

---

**Supplementary Figure S1:**
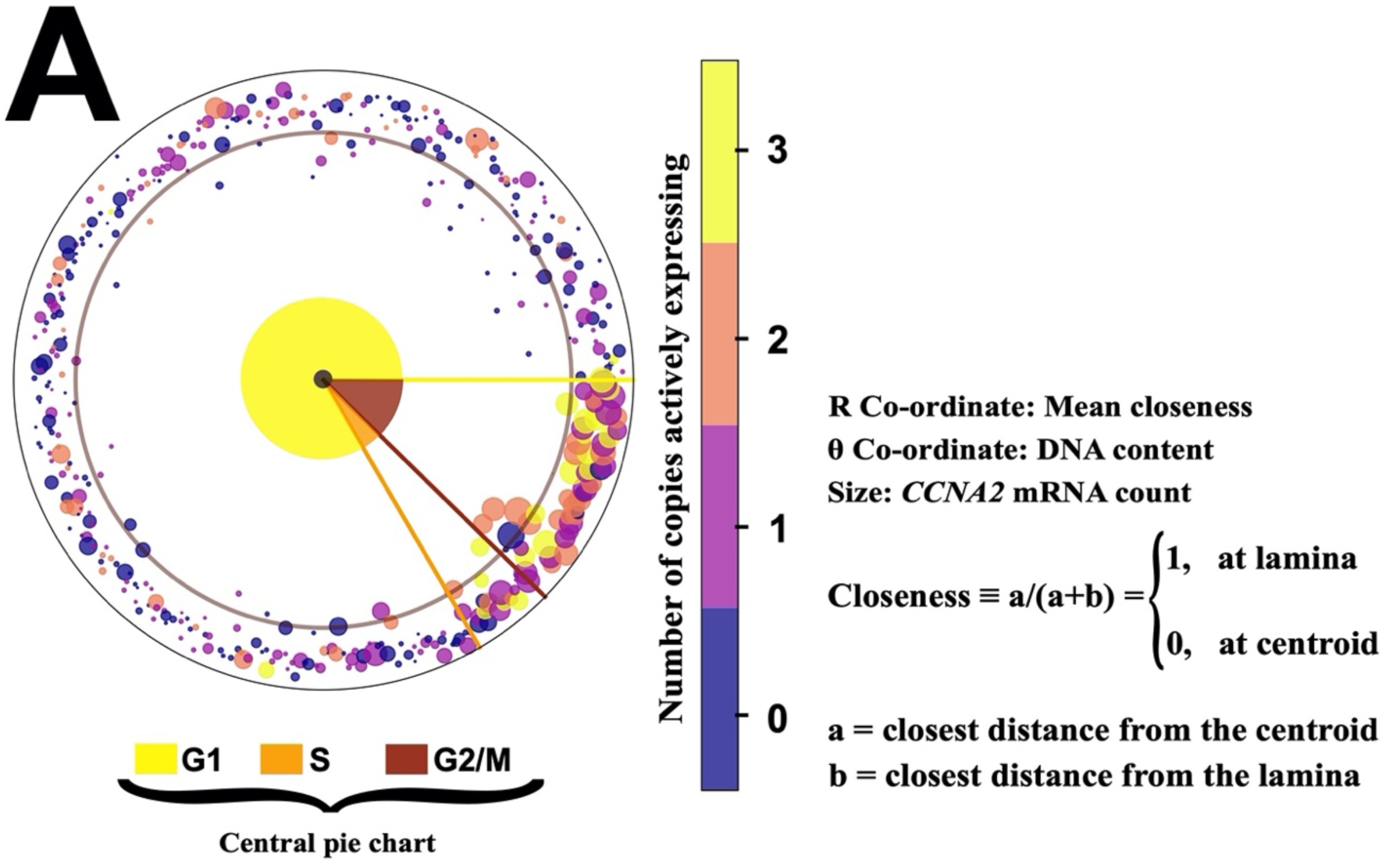
Relation between mRNA expression and the number of actively expressing copies across the cell cycle (Related to Figure 4) (A) Polar plot relating the total *CCNA2* mRNA expression with the activity of the three copies and the mean of their relative distances from the lamina in a cell. Although the number of *CCNA2* copies actively expressing increases with the increase in the *CCNA2* mRNA count of a cell in G2/M phases, there can be cells with very few mRNAs but more than a single copy expressing.

**Supplementary Table 1:**
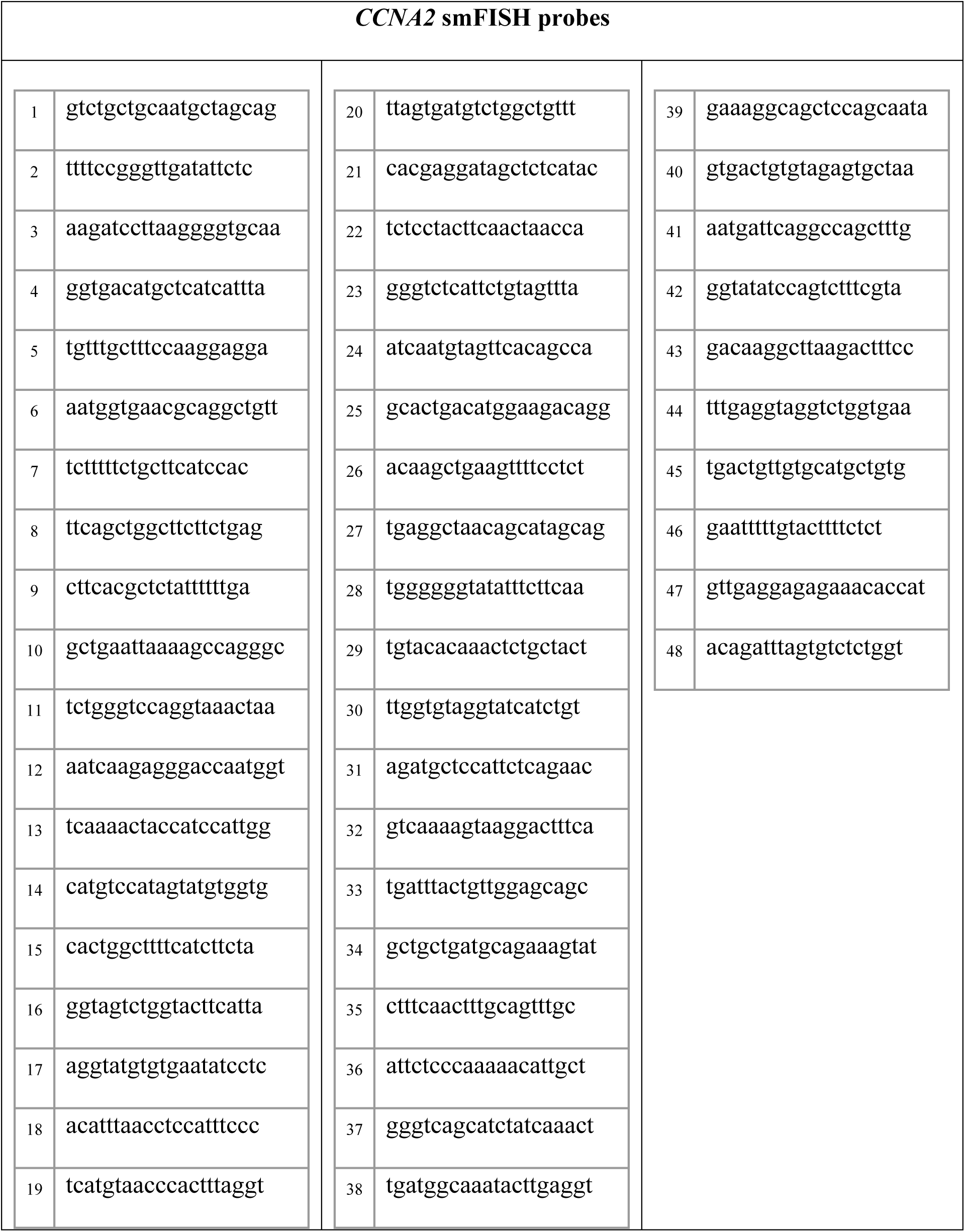
Probes Sequence (from 5’ to 3’)

